# Monocytes-derived cxcl12 guides a directional migration of blood vessels in zebrafish

**DOI:** 10.1101/2024.10.24.620141

**Authors:** Xiaofeng Lu, Xiaoning Wang, Bowen Li, Xin Wang, Xuchu Duan, Dong Liu

**Author notes:** author for correspondence Contact details for Correspondence: Dong Liu, Ph.D Seyuan Road 9, Nantong, China, 226001; Xuchu Duan, Ph.D Seyuan Road 9, Nantong, China, 226001. Xiaofeng Lu, Xiaoning Wang, and Bowen Li contributed equally to this work.

## Abstract

**Background:** Sprouting blood vessels, reaching the aimed location, and establishing the proper connections are vital for building vascular networks. Such biological processes are subject to precise molecular regulation. So far, the mechanistic insights into understanding how blood vessels grow to the correct position are limited. In particular, the guiding cues and the signaling-originating cells remain elusive.

**Methods:** Live imaging analysis was used to observe the vascular developmental process of zebrafish. Whole-mount in situ hybridization and fluorescent in situ hybridization were used to detect the expression profiles of the genes. Single-cell sequencing analysis was conducted to identify the guiding protein and its originating cells.

**Results:** Taking advantage of live imaging analysis, we described a directional blood vessel migration in the vascularization process of zebrafish pectoral fins. We demonstrated that pectoral fin vessel c (PFVc) migrated over long distances and was anastomosed with the second pair of intersegmental vessels (ISVs). Furthermore, we found the cxcl12a-cxcr4a axis specifically guided this long-distance extension of PFVc-ISV, and either inhibition or over-expression of cxcl12a-cxcr4a signaling both mislead the growth of PFVc to ectopic areas. Finally, based on an analysis of single-cell sequencing data, we revealed that a population of monocytes expresses the Cxcl12a, which guides the migration of the vascular sprout.

**Conclusions:** Our study identified Cxcl12a as the signaling molecule for orchestrating organotypic-specific long-distance migration and anastomosis of the pectoral fin vessel and ISVs in zebrafish. We discovered a specific cluster of gata1-positive monocytes that are responsible for expressing Cxcl12a. The findings offer novel insights into the mechanisms underlying organotypic vascularization in vertebrates.

## Introduction

As one of the earliest organs of vertebrates, vascular networks are essential for providing oxygen, transmitting signals to other organs, and transporting circulating metabolites and wastes [1]. A vascular network’s correct establishment is a prerequisite for its function. Misconnection of blood vessels can lead to diseases such as arteriovenous malformations [2]. Blood vessels are formed in two sequential ways: vasculogenesis and angiogenesis. While vasculogenesis mainly occurs at the early developmental stages as the de novo formation of vessels based on angioblasts, angiogenesis is a process to generate new blood vessels from preexisting ones, which involve several angiogenic events, including sprouting, elongation, anastomosis, and pruning [3–5].

In sprouting angiogenesis, anastomosis occurs when the new capillaries from parental vessels fuse with other sprouts or preexisting vessels. Anastomosis is a fundamental and essential process for building vascular networks; it is guided by tip cells and involves other cellular behaviors, including sprouting, migration, adhesion, and lumen formation, which thereby confer the anastomosis complex cellular and molecular mechanisms that are still poorly understood [6]. So far, only VE-Cadherin has been validated to be responsible for vascular fusion, and this anastomosis is caused by filopodial contacts between two proximal tip cells as an initial trigger[7]. Studies on the directional migration and connection of two distant vessels have yet to be reported, and the molecular mechanisms of blood vessel migration and anastomosis still require more investigation.

In recent years, genetic approaches have facilitated research seeking vascularization-associated functional genes. However, the studies are still hampered by the inability to observe the vascular developmental process *in vivo,* especially in mice. Zebrafish are transparent at the embryonic and juvenile stages, which allows us to observe the vascular developmental process in specific tissues and organs by live-imaging analysis[3, 8, 9]. The dorsal longitudinal anastomotic vessels (DLAV) of zebrafish formed by anastomosis of neighboring segmental arteries (SA) are particularly suitable for investigating the underlying cellular and molecular mechanisms of anastomosis[10, 11].

So far, very little has been known regarding the regulating mechanisms of directional blood vessel migration and connection, especially the involved guide cues and the signaling-originating cells. Chemokines are a family of chemotactic cytokines that play indispensable roles in vascular development. The CXCL12 (SDF-1)/CXCR4 axis has been validated to be essential for vasculature formation in diverse organs such as the kidney and gastrointestinal tract[12, 13]. Here, we used different transgenic zebrafish lines to study the organotypic vascular development in pectoral fins, which are pretty transparent during the life span. We observed the specific long-distance migration of the pectoral fin vessel and the subsequent anastomosis with the second pair of intersegmental vessels. We further validated that the monocytes-derived CXCl12a was essential for guiding this process via the CXC12a/CXCR4a axis. Our research provided new insights into the blood vessel formation during the embryonic development of zebrafish.

## Results

### Specific anastomosis between pectoral fin vessel c (PFVc) and the second pair of ISVs in the pectoral fin of zebrafish

In the previous study, Tamura *et al*. described the process of pectoral fin vessel (PFV) development in zebrafish, including that two blood vessels from the anterior and posterior sides of the fin started growing at 43 hpf and then connected to form the circumferential blood vessel loop (CBVL) at 48 hpf [14]. Here, we observed the whole angiogenic process in the pectoral fin of zebrafish embryos. We found something interesting: the PFVc grew progressively along the outer surface of the yolk towards the dorsal direction, bypassed the first pair of ISVs, and finally reached and fused with the proximal end of the second pair of ISVs at 68 hpf (Fig. 1 A-D; Supplementary Fig. 1; Movie 1). To validate the specificity of the connection between PFVc and the second pair of ISVs, timelapse imaging analysis was performed to observe the anastomotic process (Fig. 1E). The results reconfirmed the process that the PFVc grew towards the trunk and fused with the second pair of ISVs. Interestingly, during this process, although several filopodia were growing towards different directions at the beginning, only the filopodia towards the second pair of ISVs could successfully extend and fuse with the target vessels; other ones towards the other directions all degenerated and eventually disappeared (Fig. 1F-Y; Movie 2). The results indicated that the PFVc was specifically directed for long-distance anastomosis with the second pair of ISVs.

**Fig. 1.**
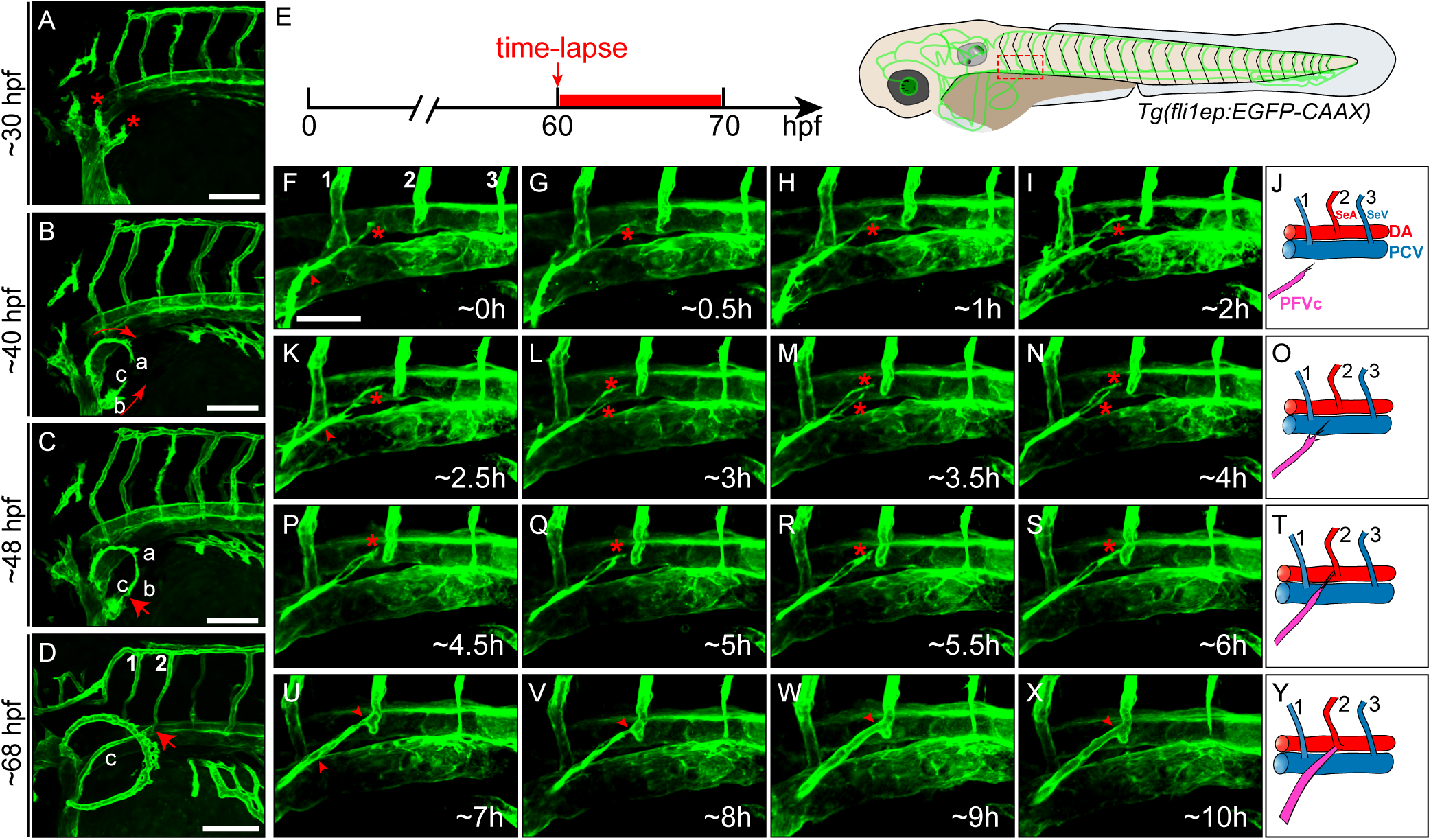
Formation of CBVL and anastomosis between PFVc and the second pair of ISVs. A, sprouts from two points of CCV towards different directions (asterisks mark the two initial sprouts from CCV. B, growth of PFVa, PFVb, and PFVc (the arrowheads show the growing directions of PFVa and PFVb). C, anastomosis between PFVa and PFVb and formation of CBVL (the arrowhead points to the anastomotic point). D, anastomosis between PFVc and the second pair of ISVs (the red arrowhead points to the anastomotic point). E-Y, Live-imaging analysis of the long-distance anastomotic process of PFVc with the second pair of ISVs on *Tg(fli1ep:EGFP-CAAX)* transgenic zebrafish. E, Schematic diagram showing the imaging time window and area. F-I, growth of PFVc towards the trunk (the asterisk marks the target ISV-directed filopodia). At the early stage, more than one filopodia sprouted from CBVL and grew in different directions (the arrowhead points to the other filopodia). K-N, the filopodia was guided to the second pair of ISVs. During this stage, only the filopodia oriented to the second pair of ISVs could extend and reach the target while the others all regressed. P-S, the connection of PA with the target ISVs. U-X, the lumenization process of PFVc-ISVs. J, O, T, Y, Schematic diagram of the anastomotic process. n=4, Scale bars: 50 μm.

### Arterial-venous identification of CBVL

To further confirm the specification of the CBVL vessel, the double transgenic zebrafish line *Tg (kdrl:Ras-mCherry::flt1:^BAC^YFP)* was employed to analyze the arterial-venous identity. The transgenic line labels the endothelial cells and arterial endothelial cells with red (*kdrl:Ras-mCherry*) and yellow (*flt1:^BAC^YFP*) fluorescence, respectively. The results showed that at the budding stage at 32 hpf, the yellow fluorescent dots were evenly distributed in PFVa, PFVb, and PFVc, indicating that these new vessels were unspecified (Fig. 2A-A’’). However, from around 40 hpf, the PFVb and PFVc started to express higher levels of *flt1* than PFVa did, indicating that PFVb and PFVc were becoming arteries, and PFVa tended to be vein (Fig. 2B-C’’). Concurrently, PFVa continuously express both flt1 and Ras from 40 hpf to 48 hpf. After the fusion between PFVc and the ISVs, and perfusion of the CBVL, green fluorescence in PFVa gradually diminished, leaving only red fluorescence at 72 hpf, indicating the venous characteristic of PFVa. Consequently, the border between the artery and vein in the CBVL was more apparent at 72 hpf (Fig. 2D-D’’). The result was further validated by using another transgenic line *Tg (lyve1b:Topza-YFP:: kdrl:RAS-mCherry),* which labels the venous cells and endothelial cells at the angiogenesis stage (Supplementary Fig. 2). In addition, we performed ISH experiment to detect the expression of *dll4, flt1* and *dab2* which specifically label the arterial and venous cells respectively. Expression of *dll4* and *flt1* could be detected at PFVb and PFVc at 48 hpf, proving both the PFVb and PFVc were arteries, while *dab2* was highly expressed at PFVa at 60 hpf, indicating the venous property of PFVa (Supplementary Fig. 3).

**Fig. 2.**
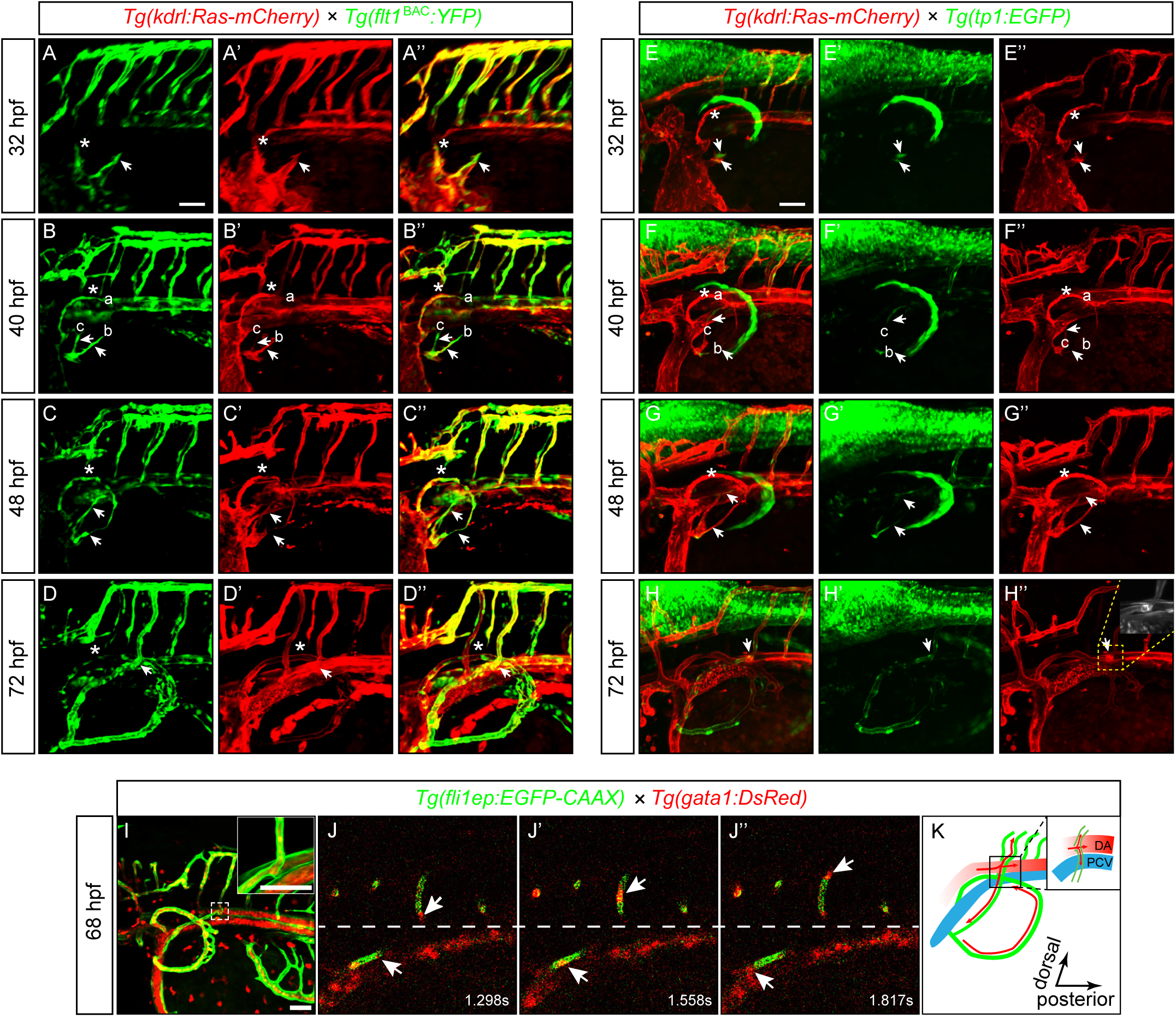
Arterial-venous specification and blood flow direction of PFVc. A-D’’, Arterial-venous specification of CBLV during the development of pectoral fin, the asterisks mark the blood vessels sprouted from the dorsal side of CCV, and the arrows mark the blood vessels sprouted from the ventral side of CCV. A-A’’, the vessels sprouted from CCV were not differentiated at 32 hpf. B-B’’, at 40 hpf, the dorsal sproutings (PFVa, marked with asterisks) grew faster than the ventral ones which branched into two new vessels (PFVb and PFVc). C-C’’, at 48 hpf, before perfusion, the PFVb and PFVc developed into an artery, while the PFVa developed into a vein indicated by the expression of *flt1* in green fluorescence. D-D’’, After perfusion, the specification of CBVL was confirmed. E-H’’, Notch signal mediated the artery development of pectoral fin vessels. E-E’’, at 32 hpf, green fluorescence was enrichedin the sporting PFVb and PFVc, but not PFVa. F-G’’, green fluorescence was observed at the tip cells of PFVb and PFVc, while no signal was detected along PFVa at 40 and 48 hpf. H-H’’, the formation and specification process of CBVL confirmed the arterial property of PFVc and the second pair of ISVs after anastomosis and perfusion of the vessels. I-J’’, The direction of blood flow in PFVc and its target ISVs. J-J’’, side view of time-lapse imaging in the white dashed box in Fig. I, arrows indicate the blood cells. K, Schematic diagram of the direction of blood flow in PFVc and its target ISVs. n=5, Scale bars: 50 μm.

Considering that Notch signaling pathway plays important roles in angiogenesis via regulating artery formation[15]. To examine the function of Notch signaling in pectoral fin vascular formation and specification, the Notch reporter line *Tg(kdrl:Ras-mCherry::tp1:EGFP)* [16] was employed to trace the vessel development and artery formation of CBVL. We observed GFP fluorescence in PFVb and PFVc since the budding stage at 32 hpf, while not in PFVa (Fig. 2E-E’’). During 40-48 hpf, GFP was continuously expressed in PFVb and PFVc, especially the area of endothelial tip cells, while not detected in PFVa (Fig. 2F-G’’). The results indicated different properties of PFVa from PFVb and PFVc. At 72 hpf, GFP fluorescence was clearly distributed in PFVb, PFVc and the second ISV (aISV), indicating their arterial property (Fig. 2H-H’’). Subsequently, we inhibited the Notch signaling by using the inhibitor LY411575, to assess the effects on vascular development. We found that the growth of PFVc was immediately inhibited when the embryos were treated by LY411575, making it unable to reach and anastomose with the target ISVs, indicating the essential role of Notch signaling in PFVc development (Supplementary Fig. 4).

In addition, we also observed that after perfusion, the erythrocytes moved from the DA to the second pair of ISVs and the PFVc, confirming the artery identity of PFVc and the second ISV (Fig. 2I-K; Movie 3). Taken together, our findings revealed the process of formation and specification of CBVL in zebrafish embryos, which suggested that the arterial-venous specification of CBVL occurred before vascular connection and perfusion.

### *cxcl12a/cxcr4a* axis regulated the directional migration of the PFVc and the anastomosis with the target ISVs

The finding that the PFVc was specifically directed to fuse with the second pair of ISVs promoted us to explore the underlying molecular mechanisms. Previous studies have revealed many molecules that guide vessel extension by regulating the vascular microenvironment or arterial-venous identification. Chemokines guide cell migration with gradients, among which Cxcl12 and Cxcl4 ligand-receptor pairs are essential for guiding endothelial migration[3]. In our initial study, single-cell transcriptome analysis of zebrafish embryonic endothelial cells at the larval stage revealed that *cxcr3.3*, *cxcr4a*, *cxcr4b*, and *cxcr7b* were expressed in vascular endothelial cells, suggesting their critical roles in regulating vascular development (Fig. 3A-E’, Supplementary Fig. 5, 6). Moreover, *cxcr4a* had a much higher expression level than *cxcr3.3, cxcr4b*, and *cxcr7b* in arterial endothelial cells, further stressing its importance in these cells (Fig. 3B-E’). Whole-mount in situ hybridization (WISH) results consistently confirmed that *cxcr4a* was specifically highly expressed in the developing CBVL (Fig. 3F-F’’). Aiming to further confirm that the directional migration of PFVc is related to *cxcr4a*, we injected AMD3100, which could competitively bind to Cxcr4a in the embryos at 48 hpf. As a result, the PFVc filopodium of treated embryos failed to reach the second pair of ISVs and eventually anastomosed with DA instead, with blood flow from DA to PFVc (Supplementary Fig. 7 and Movie 4). Furthermore, *cxcr4a* knockdown and *cxcr4a* knockout were performed. *Cxcr4a* knockdown also caused similar phenotypes with misguided PFVc, while *cxcr4a* knockout caused more serious phenotype with the loss of PFVc (Fig. 3G-K).

**Fig. 3.**
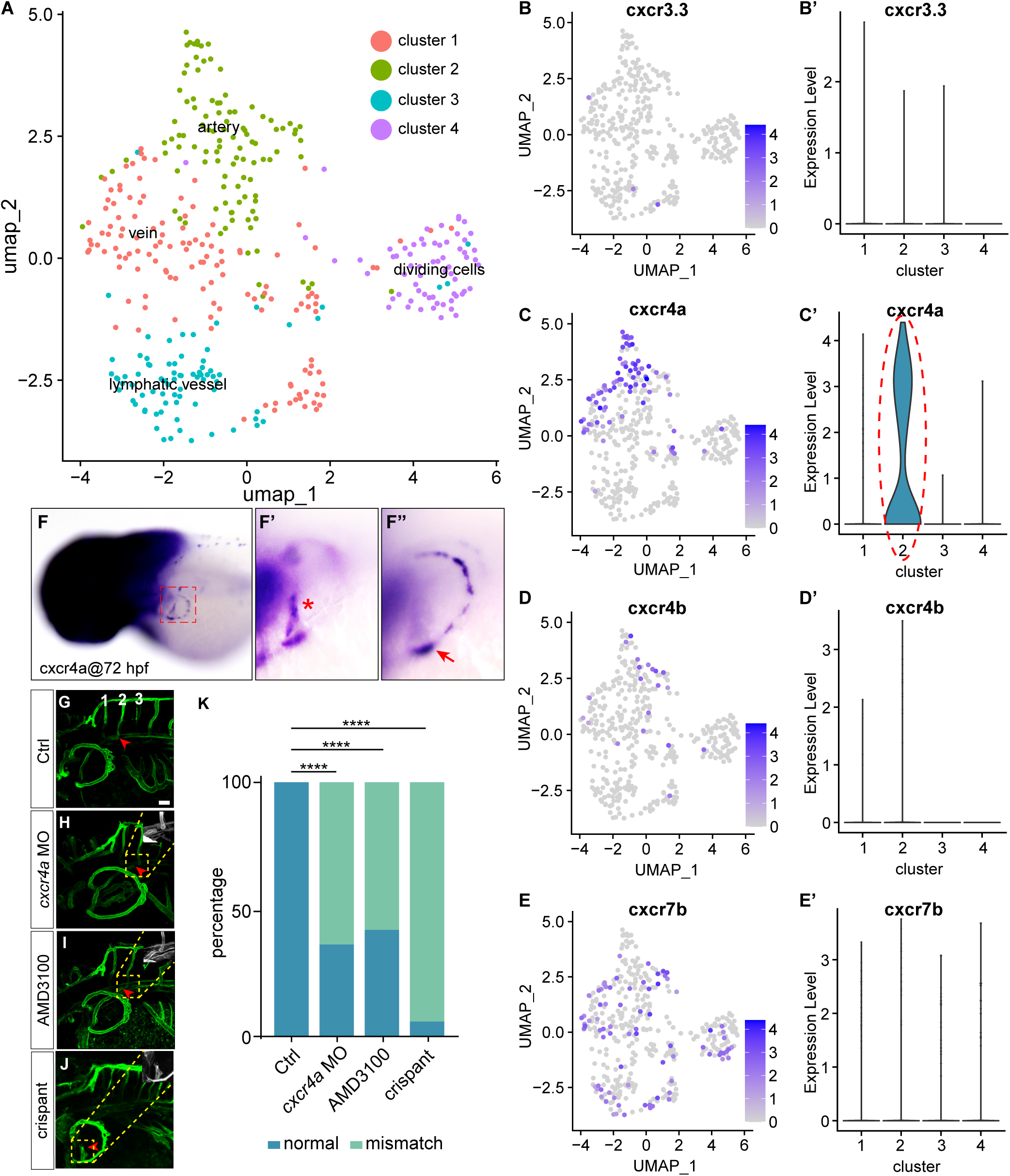
*cxcr4a* is involved in the regulation of PFVc-ISVs anastomosis. A, UMAP plot showing the clusters of fli1a positive cells (data from GSE178150). B-E’, feature plot and violin plot showing the expression of *ccxcr3.3, cxcr4a, ccxcr4b* and *ackr3b* in the four clusters in Fig. A. F-F’’, WISH analysis of *cxcr4a* expression in CBVL. G, Live-imaging analysis of the long-distance anastomosis of PFVc-ISVs on *Tg(fli1ep:EGFP-CAAX)* transgenic zebrafish. H, knockdown of *cxcr4a* by a morpholino led to the misconnection of PFVc-ISVs. I, AMD3100 treatment led to the misconnection of PFVc-ISVs. J, *cxcr4a* knockout crispant constructed by Crispr-Cas9 technique showed the loss of PFVc. K, the ratio of misconnection caused by injection of AMD3100 and knockdown of *cxcr4a*. n=20, Scale bars: 50 μm.

Since *cxcr4a* has two ligands in zebrafish, *cxcl12a* and *cxcl12b*, we examined the expression pattern of *cxcl12a* and *cxcl12b*. As expected, the result demonstrated that *cxcl12a* was highly expressed in the area around the second pair of ISVs but not the other ISVs at 72 hpf (Fig. 4A-A’’), while the *cxcl12b* was expressed in other different areas (Fig. 4B-B’). Therefore, we speculated that Cxcl12a guided the directional migration and anastomosis of PFVc to the second pair of ISVs via interacting with Cxcr4a. The subsequent experiments showed that *cxcl12a* mutation caused the misguided migration and fusion of PFVc (Fig. 4C-E). Consistently, overexpression of *cxcl12a* in larger areas of the embryos also resulted in the misconnection between PFVc and its target ISVs (Fig. 4F-H).

**Fig. 4.**
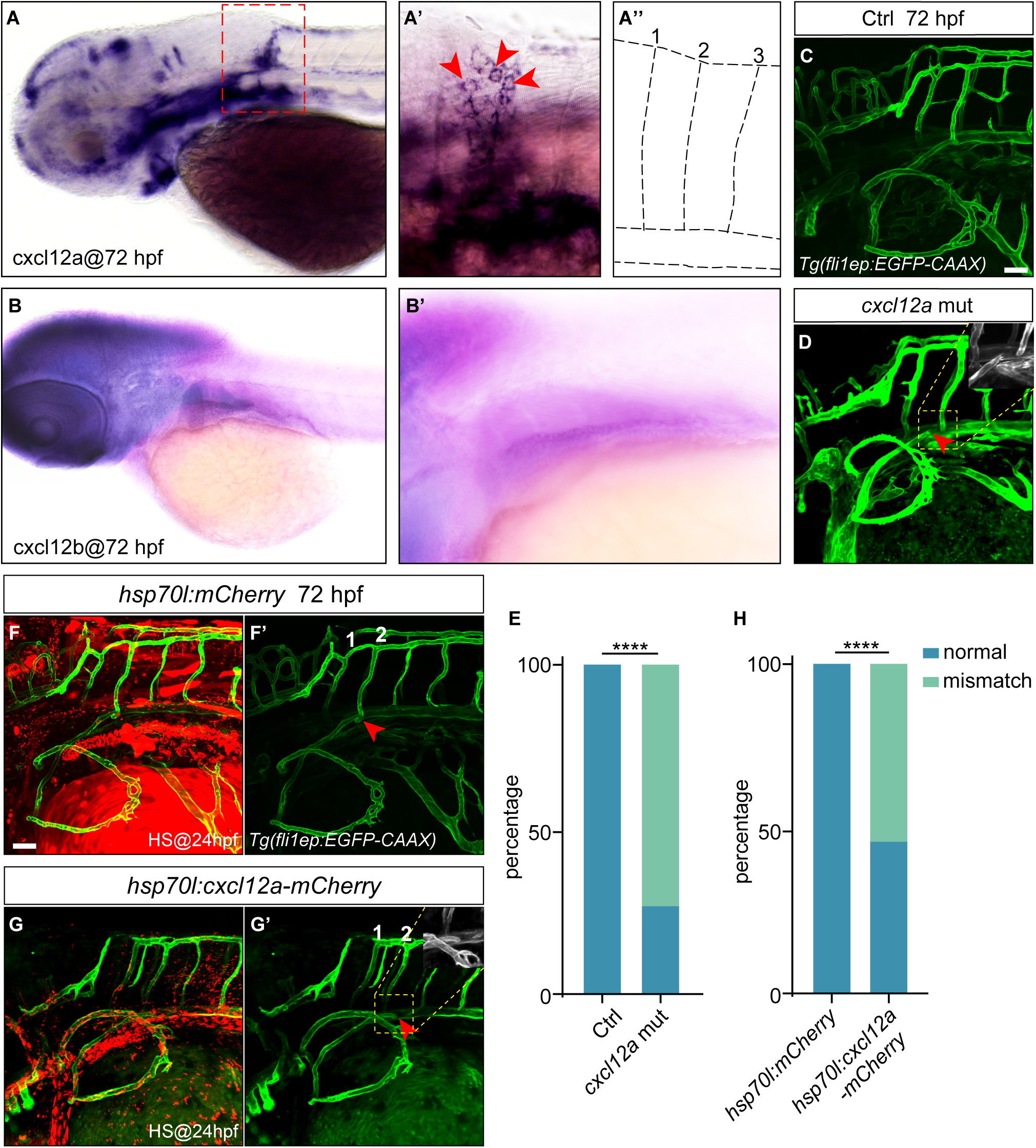
Cxcl12a is the ligand of Cxcr4a in the regulation of anastomosis of PFVc- ISVs. A-A’’, WISH analysis of *cxcl12a* expression at 72 hpf. A’, cxcl12a positive cells around the second pair of ISVs (marked with red arrowheads). A’’, the schematic illustration of the first three pairs of ISVs. B-B’, WISH analysis of *cxcl12b* expression at 72 hpf. C, Live-imaging analysis of the long-distance anastomosis of PFVc-ISVs on *Tg(fli1ep:EGFP-CAAX)* transgenic zebrafish. D, knockout of *cxcl12a* led to the misconnection of PFVc-ISVs. E, the ratio of misconnection caused by knockout of *cxcl12a*. F-F’, injection of *hsp70l:mCherry* construct did not affect the anastomosis of PFVc-ISVs. G-G’, overexpression of *cxcl12a* led to the misconnection of PFVc-ISVs. H, the ratio of misconnection caused by overexpression of *cxcl12a*. n=20, Scale bars: 50 μm.

### Cxcl12a originated from monocyte regulates PFVc growth and anastomosis

Through Whole-mount in situ hybridization (WISH), we observed that the cells expressing *cxcl12a* were morphologically similar to monocytes. To better understand these cells, we reanalyzed the single-cell RNA-sequencing data and screened the cells expressing *gata1a*, which is expressed in early myeloid cells. We found three distinct clusters (Erythrocyte 1-3), among which the Erythrocyte 2 cluster had the highest expression level and expression ratio of *cxcl12a* (Fig. 5A-C). Using the fluorescent in situ hybridization (FISH), we observed that the cells expressing *cxcl12a*, which were adjacent to the second pair of ISVs, were gata1 positive (Fig. 5D-D’’’). Subsequent analysis of the *cxcl12a* and *gata1a* co-expressing cells revealed a subset of cells within the cluster of Erythrocyte 2, which was identified as displaying relatively higher levels of co-expression of *gata1a* and *cxcl12a* (Fig. 5E-F). GO analysis showed that the genes in *gata1a-cxcl12a* positive cells were involved in locomotion, chemotaxis, blood vessel morphogenesis, and cell migration regulation, consistent with our observations (Fig. 5G). Additionally, the marker genes *pu.1*, *coro1a*, *si:ch211-212k18.7 (CD68)*, *il1b*, and *mpeg1.1*, which are indicative of monocyte and macrophage presence, were highly expressed in the subset of *gata1a-cxcl12a* co-expressing cells, suggesting that these cells may represent monocytes (Fig. 5H). To functionally prove this finding, we performed the NTR depletion of macrophage in the embryos and subsequently observed the misconnection of PFVc to DA as well (Fig. 5I-K and Movie 5,6). These findings indicate that a subset of monocytes plays a crucial role in facilitating directed long-distance vascular migration and anastomosis through the secretion of Cxcl12a (Fig. 6).

**Fig. 5.**
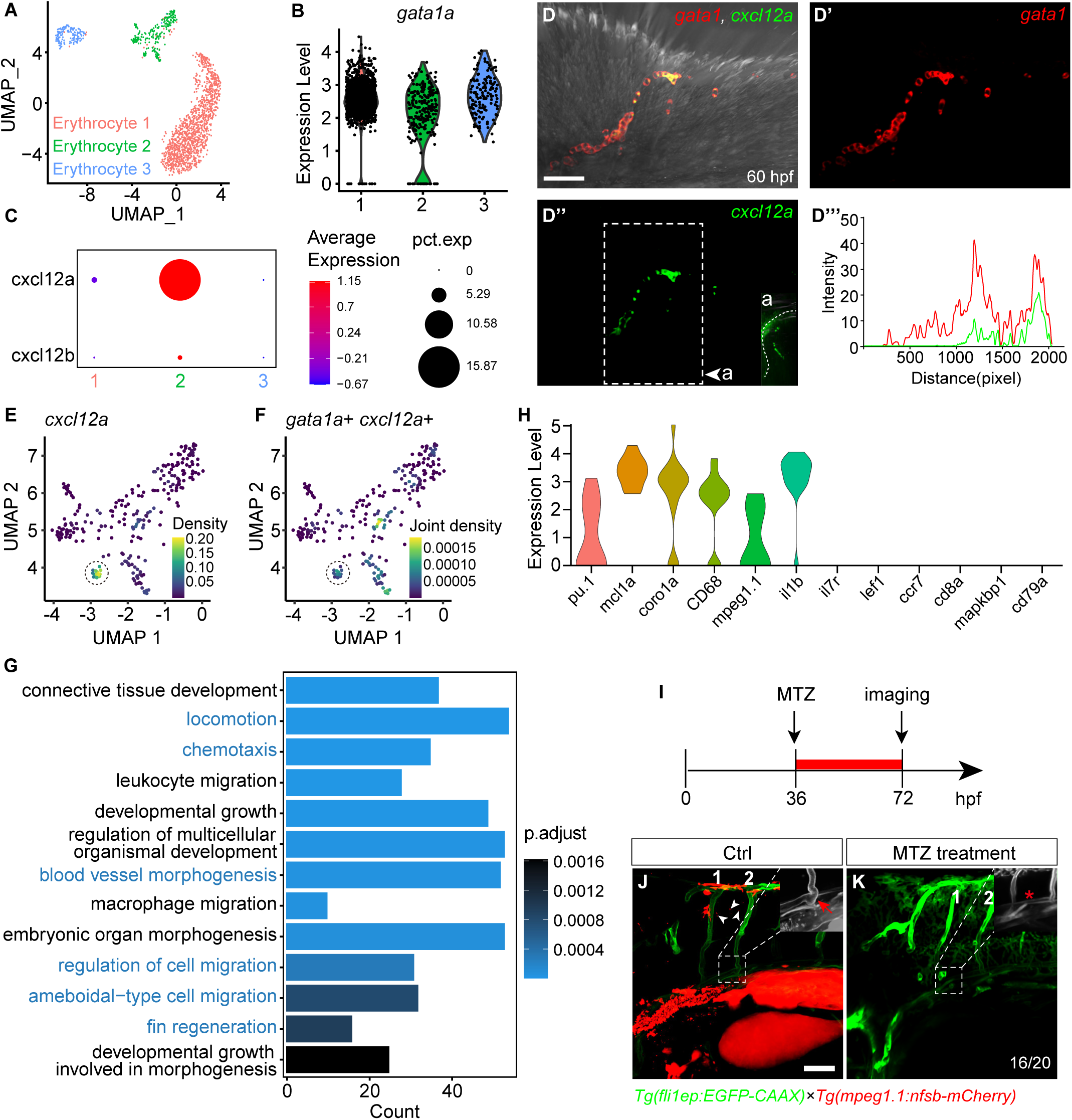
Monocytes secreted cxcl12a regulated the PFVc-ISVs anastomosis. A, UMAP plot showing the clusters of myeloid cells (data from GSE178150). B, violin plot showing the expression of *gata1a* in the three clusters in Fig. A. C, dot plot showing the average expression level and cell expression ratio of *cxcl12a* in the three clusters in Fig. A. D-D’’, FISH analysis of *cxcl12a* expression in the second pair of ISVs on *Tg (gata1: dsRed)* transgenic zebrafish embryos. D’’ side view of *cxcl12a* expression with dashed lines indicating the larva skin. D’’’, the GFP and dsRed fluorescence intensity along the second pair of ISVs in Fig. D. E-F, co-expression analysis of *gata1* and *cxcl12a* in the cells of Erythrocyte 2 cluster in Fig. A, dash circle marked the cells with highly co-expression of *gata1* and *cxcl12a*. G, GO analysis of the genes in *gata1* and *cxcl12a* co-expression cells. H, expression of the immune cell marker genes in the *gata1*-*cxcl12a* expressing cells. I, schematic diagram showing the MTZ treated NTR depletion experiment on *Tg (fli1ep: EGFP-CAAX::mpeg1.1:nfsb-mCherry)* double transgenic zebrafish. J, Live-imaging analysis of the long-distance anastomosis of PFVc-ISVs in control cluster at 72 hpf (White arrowheads indicate the mpeg1.1 positive cells and the red arrow indicates the correct anastomosis of PFVc-ISVs). K, NTR depletion of mpeg1.1 positive ccells led to the misconnection of PFVc-ISVs (marked with asterisks). n=20, Scale bars: 50 μm.

**Fig. 6.**
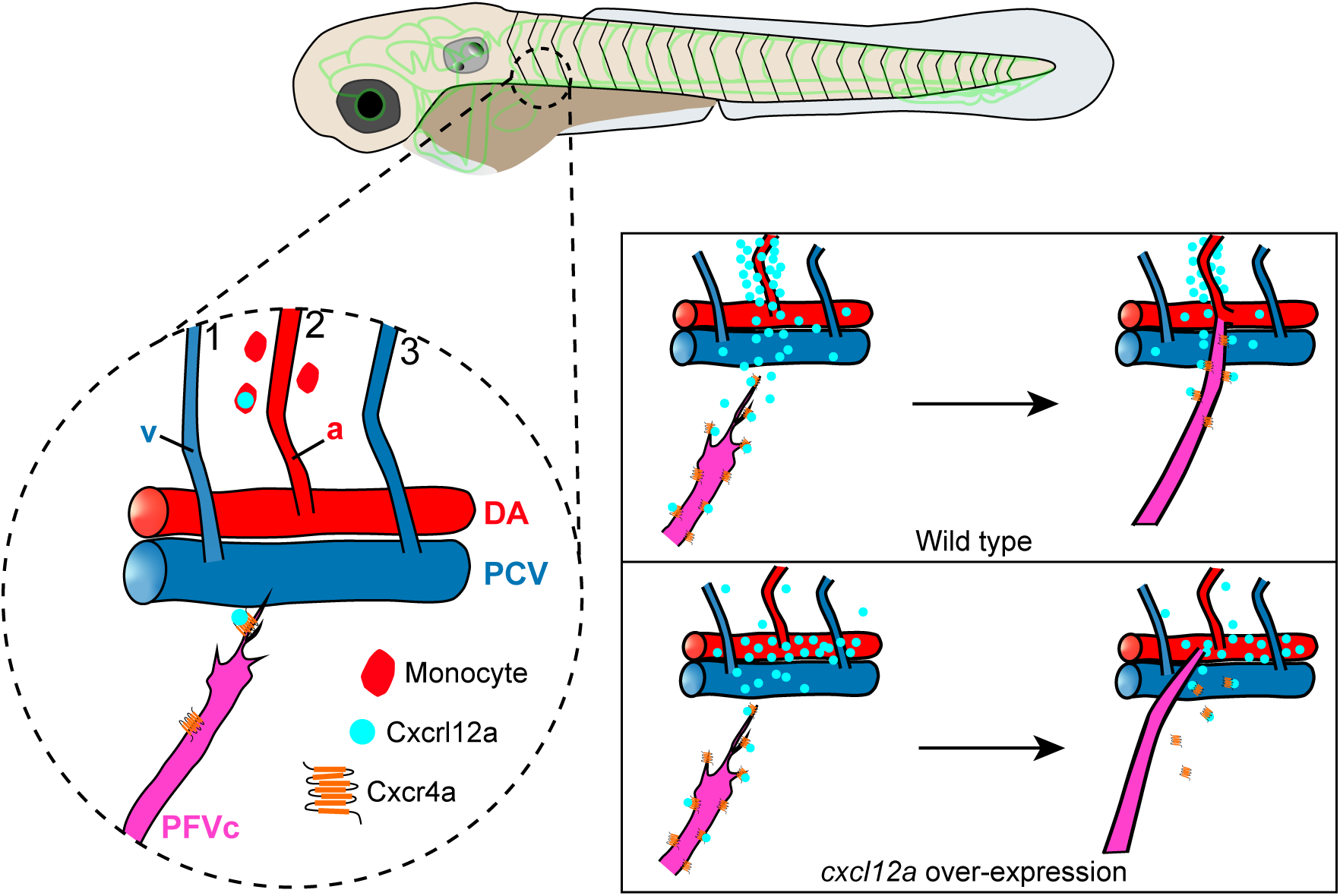
The proposed molecular mechanism guiding the directed long-distance PFVc migration and anastomosis.

## Discussion

Organotypic vasculature is essential for organs as it provides oxygen, nutrients, and signaling molecules to ensure the proper development and function of the organs. The process of organotypic vascular formation needs to be tightly coordinated with its host organ development. Although previous studies have observed the formation of the circumferential blood loop, which separates the pectoral fin into the apical fold region (AF) and the endoskeletal region, the details of the pectoral fin vascular development remain unclear[14, 17].

In this study, we elucidated the intricate molecular and cellular mechanisms underlying the anastomosis of the fin artery. Meanwhile, the PFVc extended and connected to the second pair of arterial ISVs as an artery, consistent with the previous report[18], allowing the arterial blood to move from the DA into the pectoral fin vessels and finally go to the CCV. In this respect, a question arises as to why the PFVc must establish a connection with the ISVs rather than directly with the DA. We infer that the PFVc prefers connecting to the ISVs due to their relatively equivalent vessel caliber and blood flow.

One of the fundamental questions during vascular formation is when and how arteries and veins are specified. At the beginning, the specification of the endothelial cells was proposed to be impacted by the blood flow in the vessels [19–22]. Then, studies have found that the specification began at the budding of tip cells based on genetic mechanisms before blood circulation[23–25]. So far, very few studies have been reported to elucidate the specification of organotypic vessels. In the regenerated fin, the vascularization was found to initiate from venous cells in the stump, generating new arteries and vascular plexus, which is regulated by Notch signaling[26]. Recently, another study has described the process of pectoral fin vascular formation in zebrafish[18], however, the molecular guiding cues and mechanisms involved in directing the growth and migration of the vessels remain unclear. Here, our findings demonstrated that at the early stage of sprouting before the anastomosis and perfusion of the vessels, the artery and vein have differentiated via the activation of Notch signaling directed endothelial tip cells and regulation of *flt1* expression, implying that the CBVL specification was determined genetically, which is consistent with previous study[20].

There are two kinds of vascular anastomosis during the pectoral fin development. Lenard *et al.* reported the cranial vasculature process of zebrafish embryo[27] and mentioned the two typical fusion features, including the anastomosis of PLA (palatocerebral artery) and PLA-CMV (communicating vessel). In our study, the anastomosis of PFVc-ISV was mediated by new sprouting with another preexisting vessel. Interestingly, the question of how the PFVc could specifically reach and fuse with the second pair of ISVs in such a long distance attracted our attention and prompted us to explore the mechanisms involved in this process. Considering that previous studies have shown that Cxcl12a/Cxcr4a axis mediates vascular patterns[28–30], we therefore speculated that the anastomosis between PFVc with the target ISVs might also be mediated by Cxcl12a/Cxcr4a signaling. As expected, our experiments showed highly specific expressions of *cxcr4a* and *cxcl12a* in the PFVc and around the second pair of ISVs, respectively. Besides, either binding blocking or overexpression of *cxcl12a* in other segments of zebrafish both misguided the PFVc growth, leading to further extension and fusion with other untargeted vessels (DA). Altogether, we speculate that the *cxcl12a* is expressed and distributed in gradients around the second pair of ISVs. It can bind to cxcr4a expressed on the surface of PFVc and guide the filopodium elongation from PFVc to the target ISVs. The results proved that the Cxcl12a/Cxcr4a axis was essential in mediating the long-distance specific migration and anastomosis of PFVc-ISV.

Although the role of Cxcl12a in artery development has been proved in our research and previous studies[13], the specific cellular sources responsible for its expression during this process remain unidentified. Our study has unveiled, for the first time, that a subset of monocytes containing a proportion of naïve macrophages may be implicated in the expression of *cxcl12a*. Furthermore, a recent study utilizing single-cell analysis has identified macrophages with proangiogenic activities that interact with vascular endothelial cells in various organs during prenatal development, which is in line with our result[31].

In summary, our study investigated whether the arterial-venous specification of the pectoral fin vessels was predetermined genetically. More importantly, our study revealed the specific long-distance migration and anastomosis between the pectoral fin vessel and the second pair of the ISVs and further proved that this process was mediated by the Cxcl12a/Cxcr4a axis. During this process, the *cxcl12a* was specifically expressed by monocytes. The findings provide new insights into the developmental process and molecular mechanisms of vascular formation in zebrafish.

## Methods

### Ethics statement

All animal experimentation was carried out in accordance with the NIH Guidelines for the care and use of laboratory animals (http://oacu.od.nih.gov/regs/index.htm) and ethically approved by the Administration Committee of Experimental Animals, Jiangsu Province, China (Approval ID: SYXK(SU) 2017–00121).

### Zebrafish and morpholino (MO)

Zebrafish embryos and adults were raised following previously described methods[32]. The *Tg (fli1ep: EGFP-CAAX)* zebrafish embryos were utilized to visualize the pectoral fin vessel formation process. The double transgenic *Tg (flt1:^BAC^YFP::kdrl:Ras-mCherry)* zebrafish embryos were utilized to distinguish the arterial and venous vessels. *Tg (lyve1b:Topza-YFP)* zebrafish embryos were utilized to marker venous vessels during the primary angiogenesis process. *Tg(tp1:EGFP)* zebrafish embryos labeled Notch-responsive cells. *Tg (gata1: Dsred)* zebrafish embryos were used to determine the direction of blood flow. Mo for *cxcr4* was used as previously reported [33, 34].

### Generation of genic *cxcr4a* knockout zebrafish

gRNA targeted Exon2 of cxcr4a was designed (CAGCTCTGAATTCGGCTCGG) and synthesized. The knockout zebrafish were generated by injecting *cxcr4a* gRNA (0.3 ng) with Cas9 mRNA (0.9 ng) into *Tg (fli1ep: EGFP-CAAX)* at one-cell stage embryos and then incubated in E3 solution at 28.5℃. Ten embryos were selected randomly at 24 hpf for efficiency evaluation (cxcr4a-KO-ident-F: CTTTTTCAGCACATCGTCTTTG; cxcr4a-KO-ident-R: CAGAGTGAGCACAAACAGAAGG), other embryos were transferred into PTU and incubated at 28.5℃ for imaging at 72 hpf.

### Inhibitor treatment of LY411575 and AMD3100

LY411575 is a potent γ-secretase inhibitor that targets the Notch signaling pathway. The zebrafish embryos of *Tg(tp1:GFP::kdrl:ras-mCherry)* were incubated with 2 μM and 5 μM to inhibit Notch signal from 48 hpf and 60 hpf separately, and image at 72 hpf. Treatment of AMD3100 was performed as previously reported[16].

### Whole-mount in situ hybridization (WISH) and Fluorescent in situ hybridization (FISH)

Whole-mount in situ hybridization and the preparation of RNA probes were performed as described in the previous report [35, 36]. The *cxcr4a*, *cxcl12a*, and *cxcl12b* cDNA fragments were cloned with the specific primers listed below using a wild-type embryo (AB) cDNA library. Digoxigenin-labeled antisense probes were synthesized using *an in vitro* DIG-RNA labeling transcription Kit (Roche, 11175025910) with linearized pGEM-T easy vector with each corresponding gene fragment as the templates. Zebrafish embryos and larvae were fixed with 4% paraformaldehyde (PFA) overnight at 4°C, then dehydrated with gradients of methanol and stored at −20°C in 100% methanol for subsequent analysis. The hybridization result was detected with anti-DIG-AP antibody (1:2000, Roche, 11093274910) and NBT/BCIP (1:500, Roche, 11681451001). Fluorescein RNA Labeling Mix (10×, Roche, 11685619910) was used for *in vitro* cxcl12a probe transcription for FISH.

cxcr4a-ISH-F: TTTCTCCCAACGGTGTACGG

cxcr4a-ISH-R: AGATCCATTTCTGCAGCCCC

cxcl12a-ISH-F: CCGATTTCCAACGCCAAGC

cxcl12a-ISH-R: CACGACAAACACGGAGCAAAC

cxcl12b-ISH-F: GCAATATTCGCTCTTTGGGCAAAC

cxcl12b-ISH-R: AAGGTTGGTAGGCTTAGCGG

### Generation of transgenic zebrafish over-expressing *Cxcl12a*

The coding sequence of zebrafish *cxcl12a* was amplified from the cDNA library that was established from wild-type embryos using the specific primers of *cxcl12a* F1 5’- CGGACGCGTGCCACCATGGATCTCAAAGTGATCGT-3’ and *cxcl12a* R1 5’- CGACCGGTGACCTGCTGCTGTTGGGCTT-3’, then digested with Mlu I and Age I (NEB, R3198V, R3552S) and ligated into the Hsp70-MCS-mCherry vector to generate the expression plasmid Hsp70-cxcl12a-mCherry. The transgenic zebrafish were generated by injecting hsp70: cxcl12a-mCherry plasmid (0.3 ng) with Tol2 mRNA (0.4 ng) into *Tg (fli1ep: EGFP-CAAX)* at one-cell stage embryos and then incubated in E3 solution at 28.5℃. For heat shock, embryos at 24 hpf were transferred into a 1.5 ml tube and kept in the water bath at 37℃ for 1 hour and back into the 28.5℃ to quench the heat shock.

### NTR depletion

The NTR depletion experiments were performed as described in the previous report[37]. Briefly, double transgenic *Tg (fli1ep:EGFP-CAAX::mpeg1.1:NTR-mCherry)* zebrafish embryos were used to perform the NTR depletion experiment. The embryos were transferred into the medium with 10 mM metronidazole (Sigma, M1547) at 36 hpf and imaged at 72 hpf.

### Imaging

Fluorescent imaging was performed using a Nikon A1R confocal microscope. Embryos were treated with 0.003% N-Phenylthiourea (PTU) (Sigma, P7629) at 24 hpf to suppress the skin pigmentation. Embryos were mounted in 0.7% low melting point agarose (Invitrogen, 16520100) in 1x PBS and 0.01% tricaine. All images and movies were processed and exported with Imaris (Bitplane).

For time-lapse analyses, the embryos were mounted as described above. Images were obtained every 15 minutes. Images were exported as movies using Imaris 9.0.1 (Bitplane), with four fps.

### Data availability and analysis

Whole fish scRNA-seq data at 72 hpf was obtained from the GEO database (GSE178150). Cells with specific gene expression were sorted and analyzed by tSNE or UMAP. Marker gene identification, DEG analysis, and pathway enrichment were performed. The analyses were performed based on R4.2.2.

### Statistical analysis

Statistical analyses were performed using the Prism software (Versions 8.0.2, GraphPad) with a chi-square test.

## Abbreviations

CBVL: circumferential blood vessel loop.
PFV: pectoral fin vessel.
CCV: common cardinal vein.
DA: dorsal aorta.
PCV: posterior cardinal vein
dpf: day-post-fertilization.
hpf: hour-post-fertilization.
ISV: intersegmental vessel.
SeA: intersegmental arteries.
SeV: intersegmental veins.
WISH: Whole-mount in situ hybridization.
FISH: Fluorescent in situ hybridization.

## Authors’ contributions

D Liu designed the experiments; XF Lu, XN Wang, X Wang and XC Duan performed the experiments; XF Lu, XN Wang and D Liu analyzed the data; XC Duan, XF Lu and D Liu wrote the manuscript. All authors read and approved the final manuscript.

## Availability of data and material (data transparency)

All the experimental materials generated in this study are available from the corresponding authors upon reasonable request.

## Acknowledgments

This study was supported by grants from the National Natural Science Foundation of China (2018YFA0801004 and 81870359 received by Dong Liu; http://www.nsfc.gov.cn), Natural Science Foundation for colleges and universities in Jiangsu Province (19KJB180022 received by Xuchu Duan; http://jyt.jiangsu.gov.cn/, BK2018048 received by Dong Liu), Science and Technology Foundation of Nantong City (JC22022019 received by Xiaofeng Lu; http://kjj.nantong.gov.cn/).

## Supplementary figures

**Fig. s1 Angiogenesis process of zebrafish pectoral fin.**

**Fig. s2 Arterial-venous specification of CBVL with *Tg(lyve1b:Topza-YFP::kdrl: RAS-mCherry)*.**

**Fig. s3 WISH analysis of *flt1*, *dll4* and *dab2* expression at 60 hpf.**

**Fig. s4 Notch signal inhibition causes the loss of PFVc in *Tg(kdrl:Ras-mCherry::tp1:EGFP)* zebrafish larves.**

**Fig. s5 Violin plot showing the marker genes in vein, artery, lymphatic vessel and dividing cells.**

**Fig. s6 Feature plot and violin plot showing the cxcr family genes in *fli1a* positive cells.**

**Fig. s7 Blood flow from DA to PFVc in AMD3100 treated larve with misconnected PFVc.**

## Movie Legends

**Movie 1 Process of the CBVL formation and migration of PFVc.** Time-lapse movie of the developmental process of the CBVL in Tg(fli1ep: EGFP-CAAX) embryos at 28-72 hpf. The PFVa and PFVb sprouted from CCV at about 28 hpf and formed CBVL before the anastomosis between PFVc and ISVs.

**Movie 2 The migration of PFVc and anastomosis of PFVc-ISV.** Time-lapse movie of the anastomotic process between PFVc and the target ISVs in *Tg(fli1ep: EGFP- CAAX)* embryos during 58-68 hpf.

**Movie 3 Blood flow in PFVc and the second ISVs.** The Movie shows the blood flow in the second pair of ISVs and PFVcafter the anastomosis between them., Red blood cells moved from DA to the branch of the second pair of ISVs, and then split into the second pair of ISVs and PFVc, respectively.

**Movie 4 Blood flow from DA to PFVc in mismatched larves.** In AMD3100 treated group, PFVc misconnected with DA, leading to the flow of the red blood cells from DA to PFVc.

**Movie 5, 6 The 3D view of pectoral fin vascular network of *Tg(fli1ep:EGFP- CAAX::mpeg1.1:nfsb-mCherry)* in Fig. 5J-K**. Movie 5 shows the 3D view of the connection between PFVc and the second pair of ISV in Fig. 5J. Movie 6 shows the 3D view of the connection between PFVc and DA in macrophage depleted embryos in Fig. 5K.

## References

1. Marcelo KL, Goldie LC, Hirschi KK: **Regulation of endothelial cell differentiation and specification**. Circ Res 2013, 112(9):1272–1287.

2. Schimmel K, Ali MK, Tan SY, Teng J, Do HM, Steinberg GK, Stevenson DA, Spiekerkoetter E: **Arteriovenous Malformations-Current Understanding of the Pathogenesis with Implications for Treatment**. Int J Mol Sci 2021, 22(16).

3. Schuermann A, Helker CS, Herzog W: **Angiogenesis in zebrafish**. Semin Cell Dev Biol 2014, 31:106–114.

4. Potente M, Gerhardt H, Carmeliet P: **Basic and therapeutic aspects of angiogenesis**. Cell 2011, 146(6):873–887.

5. Cunha SI, Magnusson PU, Dejana E, Lampugnani MG: **Deregulated TGF- β/BMP Signaling in Vascular Malformations**. Circ Res 2017, 121(8):981–999.

6. Charpentier MS, Conlon FL: **Cellular and molecular mechanisms underlying blood vessel lumen formation**. Bioessays 2014, 36(3):251–259.

7. Caolo V, Peacock HM, Kasaai B, Swennen G, Gordon E, Claesson-Welsh L, Post MJ, Verhamme P, Jones EAV: **Shear Stress and VE-Cadherin**. Arterioscler Thromb Vasc Biol 2018, 38(9):2174–2183.

8. Chavez MN, Aedo G, Fierro FA, Allende ML, Egana JT: **Zebrafish as an Emerging Model Organism to Study Angiogenesis in Development and Regeneration**. Front Physiol 2016, 7:56.

9. Nowak-Sliwinska P, Alitalo K, Allen E, Anisimov A, Aplin AC, Auerbach R, Augustin HG, Bates DO, van Beijnum JR, Bender RHF et al: Consensus guidelines for the use and interpretation of angiogenesis assays. Angiogenesis 2018, 21(3):425–532.

10. Herwig L, Blum Y, Krudewig A, Ellertsdottir E, Lenard A, Belting HG, Affolter M: **Distinct cellular mechanisms of blood vessel fusion in the zebrafish embryo**. Curr Biol 2011, 21(22):1942–1948.

11. Isogai S, Lawson ND, Torrealday S, Horiguchi M, Weinstein BM: **Angiogenic network formation in the developing vertebrate trunk**. Development 2003, 130(21):5281–5290.

12. Nagasawa T: Role of chemokine SDF-1/PBSF and its receptor CXCR4 in blood vessel development. Ann N Y Acad Sci 2001, 947:112–115; discussion 115-116.

13. Ara T, Tokoyoda K, Okamoto R, Koni PA, Nagasawa T: **The role of CXCL12 in the organ-specific process of artery formation**. Blood 2005, 105(8):3155–3161.

14. Yano T, Abe G, Yokoyama H, Kawakami K, Tamura K: **Mechanism of pectoral fin outgrowth in zebrafish development**. Development 2012, 139(16):2916–2925.

15. Hasan SS, Tsaryk R, Lange M, Wisniewski L, Moore JC, Lawson ND, Wojciechowska K, Schnittler H, Siekmann AF: **Endothelial Notch signalling limits angiogenesis via control of artery formation**. Nat Cell Biol 2017, 19(8):928–940.

16. Pillay LM, Mackowetzky KJ, Widen SA, Waskiewicz AJ: **Somite-Derived Retinoic Acid Regulates Zebrafish Hematopoietic Stem Cell Formation**. PLoS One 2016, 11(11):e0166040.

17. Isogai S, Horiguchi M, Weinstein BM: **The vascular anatomy of the developing zebrafish: an atlas of embryonic and early larval development**. Dev Biol 2001, 230(2):278–301.

18. Paulissen SM, Castranova DM, Krispin SM, Burns MC, Menéndez J, Torres-Vázquez J, Weinstein BM: **Anatomy and development of the pectoral fin vascular network in the zebrafish**. Development 2022, 149(5).

19. Hasan SS, Siekmann AF: **The same but different: signaling pathways in control of endothelial cell migration**. Curr Opin Cell Biol 2015, 36:86–92.

20. Pitulescu ME, Schmidt I, Giaimo BD, Antoine T, Berkenfeld F, Ferrante F, Park H, Ehling M, Biljes D, Rocha SF et al: Dll4 and Notch signalling couples sprouting angiogenesis and artery formation. Nat Cell Biol 2017, 19(8):915–927.

21. Su T, Stanley G, Sinha R, D’Amato G, Das S, Rhee S, Chang AH, Poduri A, Raftrey B, Dinh TT et al: Single-cell analysis of early progenitor cells that build coronary arteries. Nature 2018, 559(7714):356–362.

22. Fang JS, Coon BG, Gillis N, Chen Z, Qiu J, Chittenden TW, Burt JM, Schwartz MA, Hirschi KK: **Shear-induced Notch-Cx37-p27 axis arrests endothelial cell cycle to enable arterial specification**. Nat Commun 2017, 8(1):2149.

23. Moyon D, Pardanaud L, Yuan L, Bréant C, Eichmann A: **Plasticity of endothelial cells during arterial-venous differentiation in the avian embryo**. Development 2001, 128(17):3359–3370.

24. Othman-Hassan K, Patel K, Papoutsi M, Rodriguez-Niedenführ M, Christ B, Wilting J: **Arterial identity of endothelial cells is controlled by local cues**. Dev Biol 2001, 237(2):398–409.

25. le Noble F, Moyon D, Pardanaud L, Yuan L, Djonov V, Matthijsen R, Bréant C, Fleury V, Eichmann A: **Flow regulates arterial-venous differentiation in the chick embryo yolk sac**. Development 2004, 131(2):361–375.

26. Kametani Y, Chi NC, Stainier DY, Takada S: **Notch signaling regulates venous arterialization during zebrafish fin regeneration**. Genes Cells 2015, 20(5):427–438.

27. Lenard A, Ellertsdottir E, Herwig L, Krudewig A, Sauteur L, Belting HG, Affolter M: **In vivo analysis reveals a highly stereotypic morphogenetic pathway of vascular anastomosis**. Dev Cell 2013, 25(5):492–506.

28. Takabatake Y, Sugiyama T, Kohara H, Matsusaka T, Kurihara H, Koni PA, Nagasawa Y, Hamano T, Matsui I, Kawada N et al: The CXCL12 (SDF- 1)/CXCR4 axis is essential for the development of renal vasculature. J Am Soc Nephrol 2009, 20(8):1714–1723.

29. Fujita M, Cha YR, Pham VN, Sakurai A, Roman BL, Gutkind JS, Weinstein BM: **Assembly and patterning of the vascular network of the vertebrate hindbrain**. Development 2011, 138(9):1705–1715.

30. Bae YK, Kim GH, Lee JC, Seo BM, Joo KM, Lee G, Nam H: **The Significance of SDF-1alpha-CXCR4 Axis in in vivo Angiogenic Ability of Human Periodontal Ligament Stem Cells**. Mol Cells 2017, 40(6):386–392.

31. Wang Z, Wu Z, Wang H, Feng R, Wang G, Li M, Wang SY, Chen X, Su Y, Wang J et al: An immune cell atlas reveals the dynamics of human macrophage specification during prenatal development. Cell 2023, 186(20):4454–4471.e4419.

32. Wang X, Ling C, Li L, Qin Y, Qi J, Liu X, You B, Shi Y, Zhang J, Jiang Q et al: MicroRNA-10a/10b represses a novel target gene mib1 to regulate angiogenesis. Cardiovasc Res 2016, 110(1):140–150.

33. Liu J, Zhu C, Ning G, Yang L, Cao Y, Huang S, Wang Q: **Chemokine signaling links cell-cycle progression and cilia formation for left-right symmetry breaking**. PLoS Biol 2019, 17(8):e3000203.

34. Wen L, Zhang T, Wang J, Jin X, Rouf MA, Luo D, Zhu Y, Lei D, Gregersen H, Wang Y et al: The blood flow-klf6a-tagln2 axis drives vessel pruning in zebrafish by regulating endothelial cell rearrangement and actin cytoskeleton dynamics. PLoS genetics 2021, 17(7):e1009690.

35. Zhang J, Qi J, Wu S, Peng L, Shi Y, Yang J, Yin Z, Gao Y, Wang C, Gong J et al: Fatty Acid Binding Protein 11a Is Required for Brain Vessel Integrity in Zebrafish. Frontiers in physiology 2017, 8:214.

36. Huang Y, Wang X, Wang X, Xu M, Liu M, Liu D: **Nonmuscle myosin II-B (myh10) expression analysis during zebrafish embryonic development**. Gene Expr Patterns 2013, 13(7):265–270.

37. Volpe BA, Fotino TH, Steiner AB: Confocal Microscope-Based Laser Ablation and Regeneration Assay in Zebrafish Interneuromast Cells. Journal of visualized experiments : JoVE 2020(159).

